# SARS-CoV-2 infection causes transient olfactory dysfunction in mice

**DOI:** 10.1101/2020.11.10.376673

**Authors:** Qing Ye, Jia Zhou, Guan Yang, Rui-Ting Li, Qi He, Yao Zhang, Shu-Jia Wu, Qi Chen, Jia-Hui Shi, Rong-Rong Zhang, Hui-Min Zhu, Hong-Ying Qiu, Tao Zhang, Yong-Qiang Deng, Xiao-Feng Li, Ping Xu, Xiao Yang, Cheng-Feng Qin

**Author notes:** These authors contributed equally: Qing Ye, Jia Zhou, Guan Yang, Rui-Ting Li, Qi He, Yao Zhang. Correspondence (C.F.Q), (X.Y.), (P.X.).

## Abstract

Olfactory dysfunction caused by SARS-CoV-2 infection represents as one of the most predictive and common symptoms in COVID-19 patients. However, the causal link between SARS-CoV-2 infection and olfactory disorders remains lacking. Herein we demonstrate intranasal inoculation of SARS-CoV-2 induces robust viral replication in the olfactory epithelium (OE), resulting in transient olfactory dysfunction in humanized ACE2 mice. The sustentacular cells and Bowman’s gland cells in OE were identified as the major targets of SARS-CoV-2 before the invasion into olfactory sensory neurons. Remarkably, SARS-CoV-2 infection triggers cell death and immune cell infiltration, and impairs the uniformity of OE structure. Combined transcriptomic and proteomic analyses reveal the induction of antiviral and inflammatory responses, as well as the downregulation of olfactory receptors in OE from the infected animals. Overall, our mouse model recapitulates the olfactory dysfunction in COVID-19 patients, and provides critical clues to understand the physiological basis for extrapulmonary manifestations of COVID-19.

## Introduction

The Coronavirus disease 2019 (COVID-19) caused by the newly identified severe acute respiratory syndrome coronavirus 2 (SARS-CoV-2) has caused global crisis. The clinical manifestations caused by SARS-CoV-2 predominantly involves the respiratory system, including cough, sore throat, pneumonia, and acute respiratory distress syndrome (ARDS) (**Huang et al., 2020**; **Wang et al., 2020**). With the wide spreading of the disease, a significant portion of COVID-19 patients developed anosmia, hyposmia or other olfactory dysfunctions according to clinical reports (**Giacomelli et al., 2020**; **Menni et al., 2020**; **Wölfel et al., 2020**). Accumulated evidence has established the alteration of smell as one of the most predictive symptoms for COVID-19 screening (**Menni et al., 2020**; **Spinato et al., 2020**).

The perception of smell begins with the odorant binding to the olfactory receptors (ORs) of olfactory sensory neurons (OSNs) along the upper surface of olfactory epithelium (OE). Each OSN projects an axon into the glomerulus of the olfactory bulb (OB) and then synapses with the second order neuron to convey the odor information into the olfactory cortex. Previously, upper respiratory tract infections have been considered as a common cause of olfactory disorders. Mouse models have been used to reproduce the olfactory infection and subsequent dysfunction (**Kobayakawa et al., 2007**; **Papes et al., 2018**). For example, the post viral olfactory disorders was observed in Sendai virus infected mice by buried food pellet test (BFPT), as well as the impairment of OE and OB tissues (**Matsunami et al., 2016**). However, the animal model that can recapitulate the olfactory dysfunctions seen in COVID-19 patients has not been established to date.

Human nasal respiratory epithelium (RE) cells possess an enriched expression of angiotensin-converting enzyme 2 (ACE2) (**Sungnak et al., 2020**; **Ziegler et al., 2020**), the functional receptor of SARS-CoV-2 (**Hoffmann et al., 2020**; **Walls et al., 2020**; **Zhou et al., 2020**). Single-cell RNA sequencing analyses have characterized the expression profile of ACE2 in the OE of mouse and human, mainly in non-neuroepithelium cells (**Brann et al., 2020**; **Ziegler et al., 2020**), and a recent study based on hamster model has also observed plenty of SARS-CoV-2 infected cells in the OE section (**Bryche et al., 2020**; **Sia et al., 2020**). Besides, vascular pericytes in OB were validated to possess a high level expression of ACE2 in mouse model (**Brann et al., 2020**), which play a key role on the maintenance of blood-brain barrier, as well as the regulation of blood pressure and host immune response (**Armulik et al., 2011**). Interestingly, some respiratory viruses, such as influenza virus, respiratory syncytial virus, are able to invade the OB and other parts of brain to establish infection (**Dubé et al., 2018**; **Netland et al., 2008**). Thus, how SARS-CoV-2 invade the olfactory system and contribute to the observed central nervous system (CNS) diseases remains to be determined. In the present study, we demonstrate that SARS-CoV-2 infection directly cause transient olfactory dysfunction in an established mouse model, and characterized the major target cells and pathological effects attributed to the olfactory dysfunction.

## Results

### SARS-CoV-2 targets OE and causes transient olfactory dysfunction in hACE2 mice

We have previously established a humanized ACE2 (hACE2) mouse model susceptible to SARS-CoV-2 infection (**Sun et al., 2020**). Herein, to determine the impact of SARS-CoV-2 infection on olfactory system, groups of 6-8 weeks old hACE2 mice were intranasally infected with 5.4 × 10^5^ plaque-forming units (PFU) of SARS-CoV-2. Mice inoculated with the same volume of culture media were set as mock infection controls. At 2- and 4-days post infection (dpi), tissues from the respiratory tract and olfactory system were collected from the necropsied mice, respectively, and subjected to virological and immunological assays (**Figure 1A**). As expected, high levels of SARS-CoV-2 RNAs were detected in the nasal respiratory epithelium (RE), trachea and lung at 2 and 4 dpi, and peak viral RNA (2.36×10^11^ RNA copies/mouse) was detected in the lung at 2 dpi (**Figure S1A**). Robust viral nucleocapsid (N) protein was detected in the lung from SARS-CoV-2 infected hACE2 mice, but not from the control animals (**Figure S1B**). Strikingly, high levels of viral RNAs (5.85×10^9^ RNA copies/mouse) were also detected in the olfactory mucosa (OM) at 2 dpi and maintained at high level (8.93×10^8^ RNA copies/mouse) till 4 dpi (**Figure 1B**), while the viral RNA levels were much lower in the OB and other parts of brain on 2 dpi and decreased to marginal level on 4 dpi. Furthermore, immunofluorescence staining assay detected a large amount of SARS-CoV-2 N proteins in the OE along OM (**Figure 1C**), while no viral N protein was detected in the OB and other parts of brain from SARS-CoV-2 infected hACE2 mice (**Figure S1C**). Additionally, *in situ* hybridization (ISH) by RNAscope demonstrated that SARS-CoV-2 RNA was predominantly detected in the OE (**Figure S1D**), but no in the OB (**Figure S1E**).

**Figure 1.**
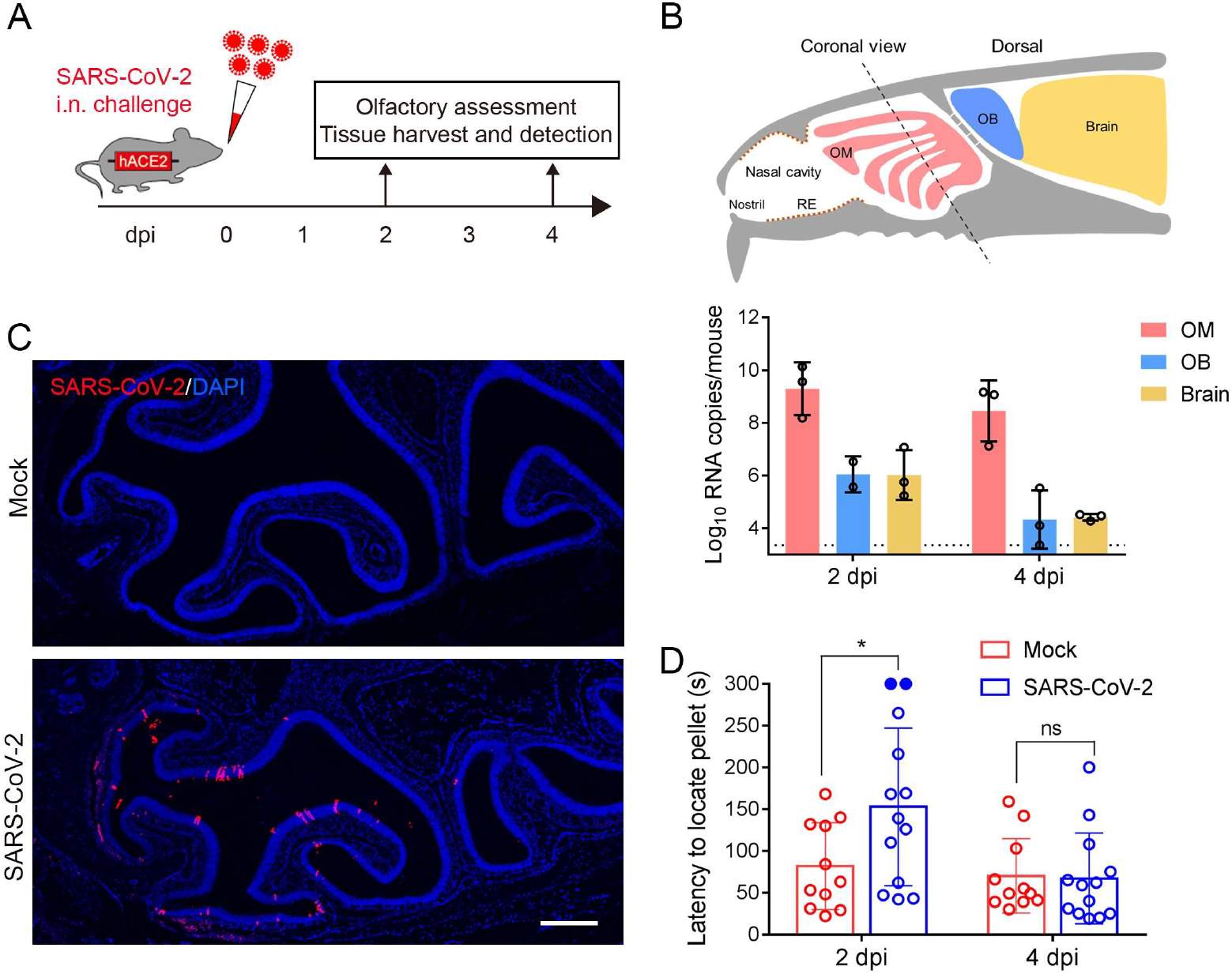
SARS-CoV-2 primarily infects the OE and causes olfactory dysfunction in hACE2 mice. (A) Schematic diagram of experimental design. Briefly, groups of 6-8 weeks old hACE2 mice were infected with 5.4 × 10^5^ PFU of SARS-CoV-2 intranasally. Olfactory function of infected mice was measured by the buried food pellet test at indicated times post inoculation. Mice were sacrificed at 2 dpi and 4 dpi for viral detection and histopathological analysis. (B) Schematic view of the OM in the nasal cavity of mice in a sagittal plane, the dotted line indicated a coronal section (upper). And viral RNA copies were determined by real-time qPCR and shown as mean ± SD from three independent replicates (lower). (C) Immunostaining of OM from SARS-CoV-2 infected mice for SARS-CoV-2 N protein (red) and DAPI (blue). Scale bar, 400 μm. (D) Buried food pellet test. Latency to locate the food pellets for mice infected with SARS-CoV-2 (n=13) or DMEM (n=11) was measured at 2 dpi and 4 dpi. See also Figure S1.

To examine whether SARS-CoV-2 infection directly impairs the olfactory function of infected mice, a standard BFPT was conducted on 2 and 4 dpi, respectively. Remarkably, a significantly increased latency (152.8 s *v.s*. 81.8 s; *p*=0.022) to locate food pellets was observed in SARS-CoV-2 infected mice as compared with the control animals on 2 dpi (**Figure 1D**). Of particular note, 2 out of 13 infected mice developed severe symptoms of anosmia as they failed to locate the food pellet within the observation period. Interestingly, recovery from olfactory dysfunction of infected mice was observed at 4 dpi, as the latency to locate food pellets was no difference from that of the control animals (67.1 s *v.s.* 70.2 s; *p*=0.992). Thus, these results demonstrate that SARS-CoV-2 primarily infects OE and leads to olfactory dysfunction in mice.

### SARS-CoV-2 primary targets non-neuroepithelial cells in the OE of hACE2 mice

The OM consists of OE and the underlying lamina propria (LP). The OE is composed of olfactory stem/progenitor cells including the horizontal basal cells (HBCs) and globose basal cells (GBCs) residing in the basal region, the mature and immature OSNs, and a variety of non-neuroepithelial lineage including the sustentacular cells, microvillar cells and Bowman’s gland cells. The OSNs lining under the supporting cells project numerous dendritic cilia with ORs into the nasal cavity and intermingle with the microvilli of sustentacular cells and microvillar cells (**Figure S2A**). Due to the asymmetrical expression pattern of ACE2 on the cell membrane as well as the unique organization of OE, it is not easy to determine which cell compartments express ACE2.

To overcome this, we took advantage of the tdTomato cassette downstream of hACE2 transgene with an internal ribosome entry site (IRES), which allows the detection of hACE2 expression by cytoplasmic fluorescence of tdTomato (**Figure S2B**). An abundant expression of hACE2 along the apical surface of OE as well as within the underlying LP was detected with a human ACE2-specific monoclonal antibody, exhibiting a similar expression pattern of tdTomato (**Figure S2C**). A detailed characterization of hACE2/tdTomato expressing cells in OM revealed that non-neuroepithelial cells, including the sustentacular cells (CK8-postive, **Figure S2D, d1**), the duct and acinus of Bowman’s gland cells (Sox9/CK8-positive, **Figure S2D, d2, d4**) in the OE and LP, respectively, and the microvillar cells (CD73/CK8-positive, **Figure S2E**), are the primary cell types that harbor human ACE2 expression (**Figure S2D**), whereas little hACE2/tdTomato expression was detected in the neuroepithelial lineage, including HBCs (CK5-positive), GBCs (Sox2-positive at the basal region), immature olfactory sensory neurons (iOSNs) (GAP43-positive) and mature olfactory sensory neurons (mOSNs) (OMP-positive) (**Figure S2D, d1-d4**).

To further characterize the primary targets of SARS-CoV-2 in the OE, multiplex immunostaining assays were performed with antibodies against SARS-CoV-2 and specific cell markers. Remarkably, robust expression of SARS-CoV-2 viral N protein was detected in the non-neuroepithelial lineage lining the outer surface of OE at 2 and 4 dpi (**Figures 2A and 2C**). The sustentacular cells (58.97%) and Bowman’s gland cells (22.76%) represent as the major target cell types at 2 dpi, while some microvillar cells (6.93%) and HBCs (4.11%) were also infected by SARS-CoV-2 (**Figures 2A and 2B**). Additionally, a small population of iOSNs (1.28%) were also infected by SARS-CoV-2, while none mOSN was infected at 2 dpi (**Figures 2A and 2B**). Interestingly, SARS-CoV-2-positive HBCs and iOSNs were found adjacent to infected sustentacular cells (**Fig. 2a**). Additionally, substantial viral protein was detected within the cilia, the cellular bodies and the underlying nerve bundles of mOSNs at 4 dpi (**Figure 2C, c1-c2**). These results indicated that SARS-CoV-2 primarily targets the non-neuroepithelial cells lining the outer surface of OE, and subsequently invades the neuroepithelial lineage in hACE2 mice.

**Figure 2.**
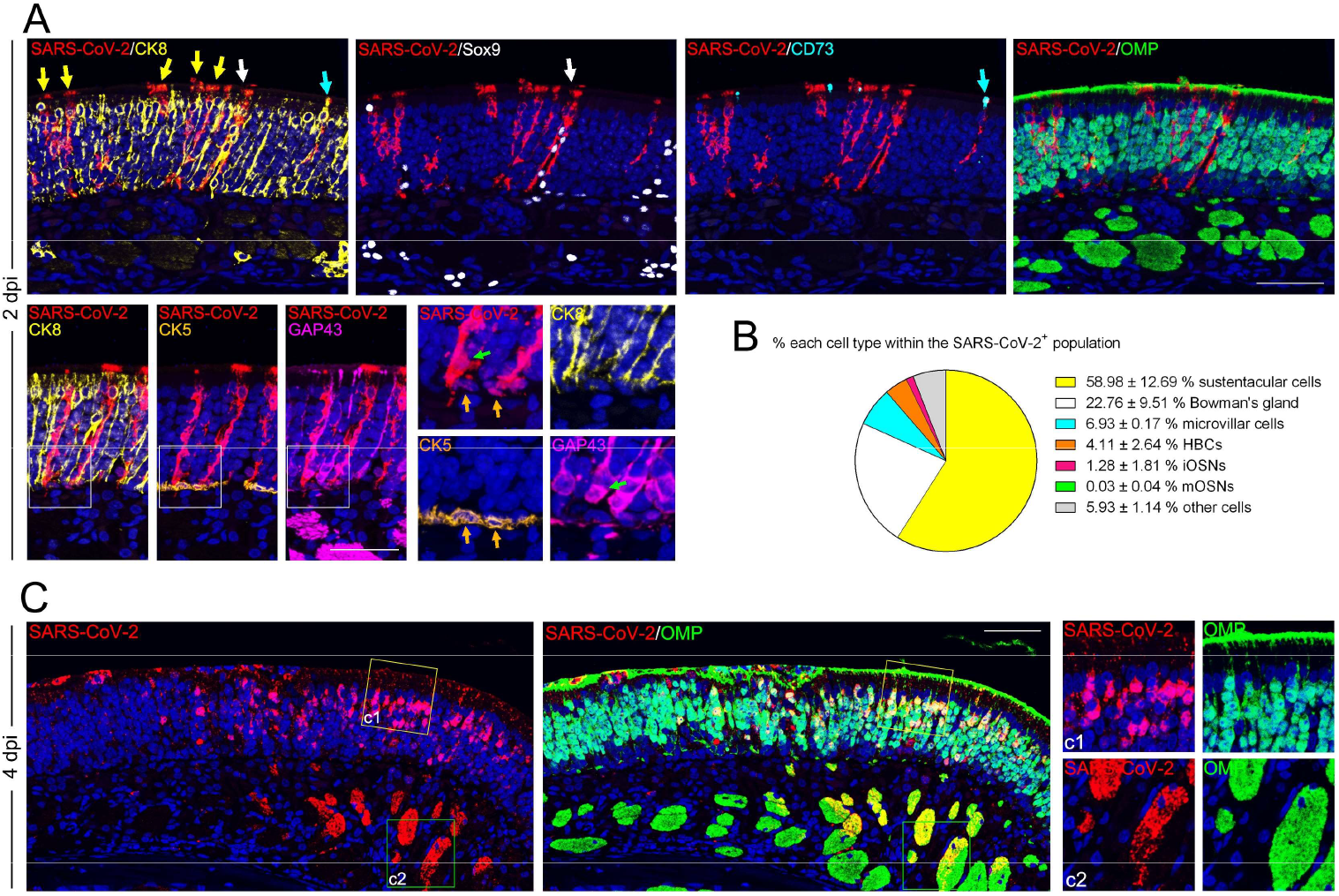
SARS-CoV-2 primarily targets non-neuroepithelial cells in the OE. (A) Representative multiplex immunofluorescent staining shows SARS-CoV-2 (SARS-CoV-2 N protein-positive) infects sustentacular cells (CK8-positive, yellow arrows), Bowman’s gland cells (Sox9/CK8-positive, white arrows), microvillar cells (CD73/CK8-positive, cyan arrows), HBCs (CK5-postitive, gold arrows) and iOSNs (GAP43-positive, green arrows) at 2 dpi. Little SARS-CoV-2 N protein is detected within OMP-positive mOSNs. (B) Statistical analysis of the percentage of each cell compartment within the SARS-CoV-2-positive cells. Data were presented as mean ± SD (n = 3). (C) Multiplex immunofluorescent staining shows an OM sample at 4 dpi with SARS-CoV-2 detected in the OMP-positive mOSNs and the underlying nerve bundles. The framed areas labelled as c1 and c2 are shown adjacently at larger magnifications. Scale bar, 50 μm. See also Figure S2.

### SARS-CoV-2 infection triggers apoptosis and immune cell infiltration in OE

We then characterized the histopathological changes of OE in response to SARS-CoV-2 infection. Strikingly, SARS-CoV-2 infection directly impaired the structural uniformity of OE, as characterized by clusters of remnants on the surface of OE (**Figure 3A**), as well as disorganized arrangement of supporting cells (**Figure 3B**) and olfactory neurons (**Figure 3C**). The integrity of the cilia layer of mOSNs and the microvilli of supporting cells were severely damaged (**Figures 3B and 3C**). More importantly, compared with mock treated groups, profound cell apoptosis (cleaved-caspase3-positive) was observed in both of the OE and LP section of OM from the SARS-CoV-2 infected mice (**Figure 3D**). Immunofluorescence co-staining indicated the apoptosis can be seen in sustentacular cells, HBCs as well as the cellular bodies and the underlying nerve bundles of iOSNs and mOSNs (**Figure 3D**). Additionally, the infiltrations of immune cells, including the macrophages (CD68-positive), the dendritic cells (CD103-positive) and the neutrophils (Ly-6G-positive) were evident in the infected OE (**Figure 3E**). The profound invasion of CD8 T lymphocytes with high expression of cytotoxic enzymes Perforin and Granzyme B would further deteriorate the cellularity of olfactory epithelial cells (**Figure 3F**). These observed physiological damages upon to SARS-CoV-2 infection probably contribute to the functional loss of olfaction.

**Figure 3.**
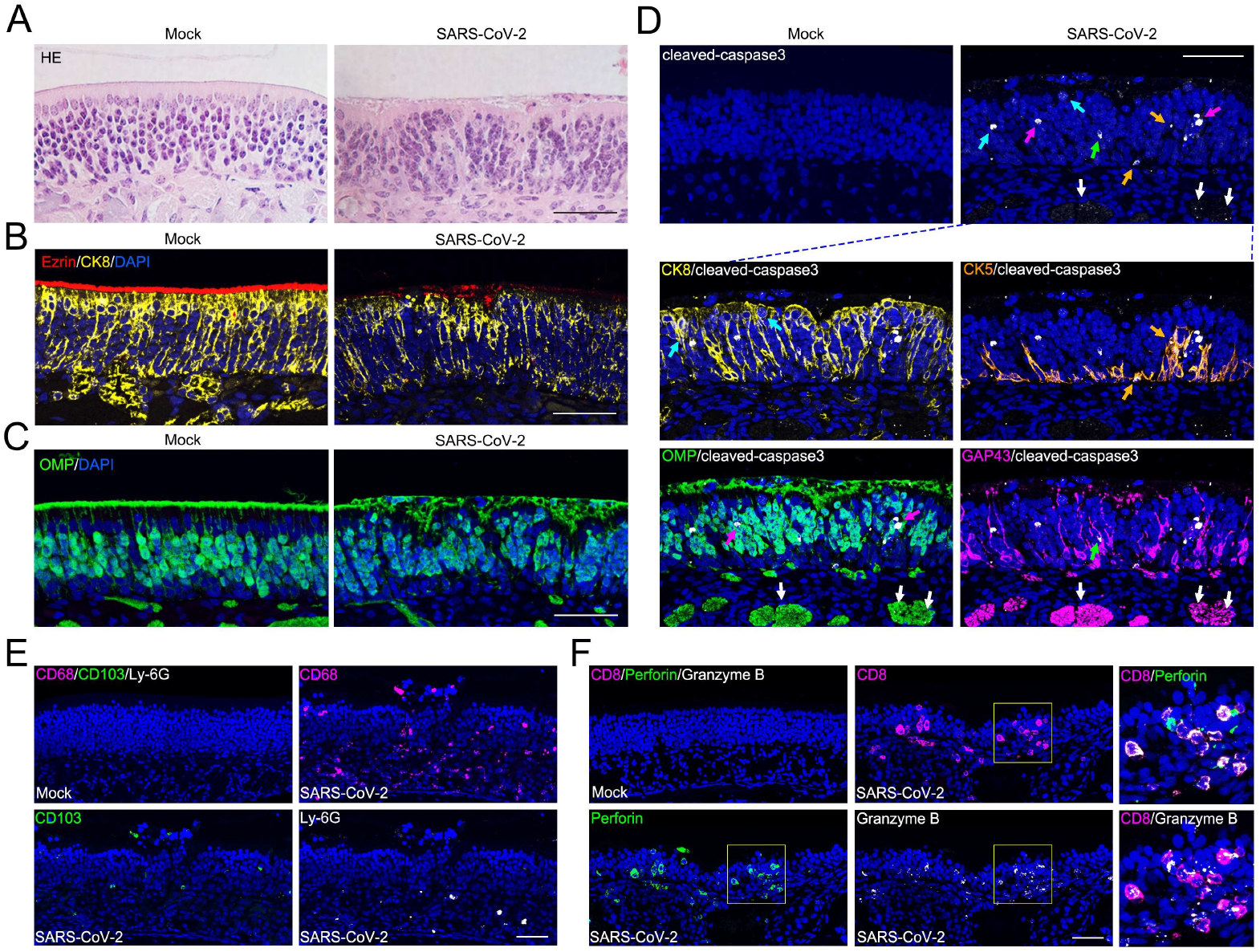
SARS-CoV-2 infection induces apoptosis and immune cell infiltration in OE. (A) Representative hematoxylin-eosin (HE) shows histopathological changes of OE. (B) Representative multiplex immunofluorescent detection of sustentacular cells (CK8-positive) and microvilli (Ezrin-positive) of OE. (C) Representative immunofluorescent detection of mOSNs (OMP-positive) of OE. (D) Apoptosis of olfactory epithelial cells (cleaved-caspase3-positive, white) after SARS-CoV-2 infection. The panels below shows apoptosis of sustentacular cells (CK8-positive, yellow; indicated by cyan arrows), HBCs (CK5-positive, gold; indicated by gold arrows), mOSN (OMP-positive, green; indicated by magenta arrows), iOSN (GAP43-positive, magenta; indicated by magenta arrows) and olfactory nerve bundles (OMP/GAP43-positive; indicated by white arrows). (E) Representative multiplex immunofluorescent staining shows infiltration of macrophages (CD68-positive, magenta), dendritic cells (CD103-positive, green) and neutrophils (Ly-6G-positive, white) in the OE after infection. (F) Representative multiplex immunofluorescent staining shows infiltration of CD8 cytotoxic T lymphocytes (magenta) with expression of Perforin (green) and Granzyme B (white) in the olfactory mucosa after infection. The framed areas are shown adjacently at larger magnifications. Scale bar, 50 μm.

### SARS-CoV-2 infection induces regeneration of OE

Without infection, HBCs at the basal region of OE remains quiescent as indicated by little expression of the proliferation marker Ki67 within CK5-positive cells (**Figure 4A, a1**). SARS-CoV-2 infection significantly increased the number of CK5/Sox2/Ki67 triple-positive cells, strongly suggesting a transition from HBCs to actively cycling GBCs (**Figure 4A, a2**). Of particular note, a prominent upward growth of HBCs from the basal layer into the upper section of OE was observed in infected animals, which also co-express the markers of their lineage offspring such as iOSNs (**Figure 4B, b1**), sustentacular cells (**Figure 4B, b2**) and the microvillar cells (**Figure 4B, b3**). These results suggest that the impaired OE is regenerated through olfactory stem cell-based proliferation and differentiation into olfactory neurons and supporting lineage, thereby restoring the normal function of OE.

**Figure 4.**
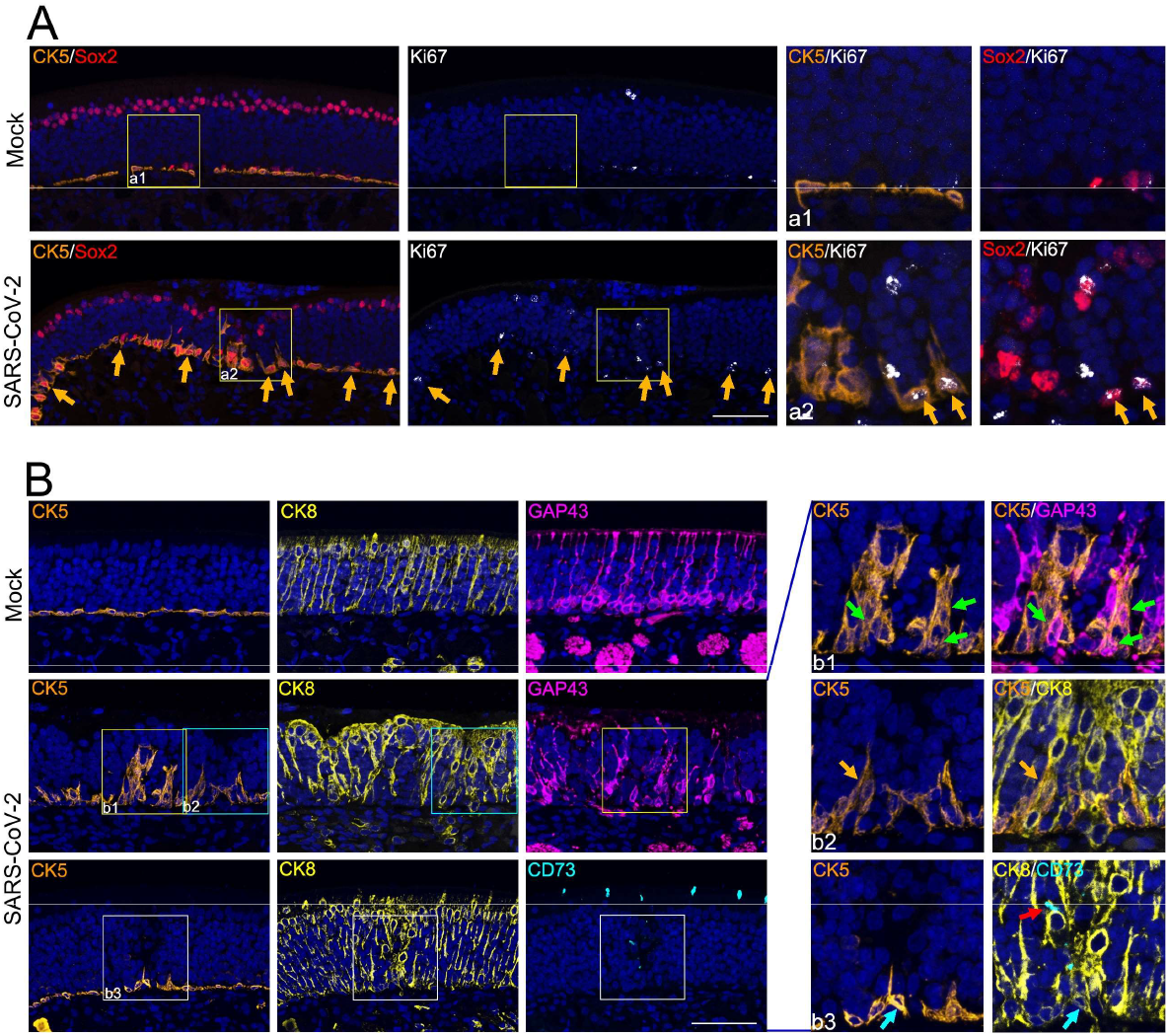
SARS-CoV-2 infection triggers regeneration of OE. (A) Representative immunofluorescent staining of CK5 (gold), Sox2 (red) and Ki67 (white) shows the increase of actively cycling olfactory stem cells as labelled CK5/Sox2/Ki67-triple-positive after infection (gold arrows). The framed areas labelled as a1 and a2 are shown adjacently at larger magnifications. (B) Representative immunofluorescent staining of CK5 (gold), CK8 (yellow), CD73 (cyan) and GAP43 (magenta) shows the transition states during the differentiation of HBCs. The framed areas labelled as b1–b3 are shown adjacently at larger magnifications. Green arrows in b1 denote CK5/GAP43 double-positive cells. Gold arrows in b2 denote CK5/CK8 double-positive cells. Cyan arrows and red arrow in b3 denote CK5/CK8 and CK8/CD73 double-positive cells, respectively. Scale bar, 50 μm.

### SARS-CoV-2 infection induces inflammatory response and suppresses olfactory signaling pathway in OE

To decipher the underlying mechanism of the observed olfactory dysfunction in SARS-CoV-2 infected mice at the molecular level, combined transcriptomic and quantitative proteomic analyses of the OE and OB samples from SARS-CoV-2 infected mice were performed in comparison with that from the control animals. In the OE samples, a total of 939 genes and 507 proteins were regulated upon SARS-CoV-2 infection, and 40 of them were synchronously regulated at both mRNA and protein levels (**Figures S3A and S3B**). While in the OB samples, 286 genes and 251 proteins were up/down regulated, and only 4 of them were consistently regulated at mRNA and protein levels (**Figures S3A and S3C**). These results further support that OE represents the major site for SARS-CoV-2 replication. Gene enrichment analyses showed that SARS-CoV-2 infection induces strong antiviral defense and inflammatory response in OE at both mRNA and protein levels at 2 dpi, which faded at 4 dpi (**Figures 5A, S4A and S4C**). Moreover, genes related to “positive regulation of cell death” and “regulation of neuron projection development” were also up regulated upon SARS-CoV-2 infection (**Figures 5A, S4B and S4D**), which was consistent with the immunostaining results (**Figure 3D and 4A**). Further integrated omics analysis of the OE samples showed that a total of 30 genes were up regulated at both mRNA and protein levels. Of which, antiviral genes/proteins including Isg15, Stat1, Stat2, Oasl2, Ifit2, Ifit3, etc, were found to interact closely. Other genes/proteins involved in neurotransmitter transport including Erc2, Lin7a, Slc1a3 and Slc25a18, were also observed (**Figure 5B**). While in the OB samples, we did not find any induction of antiviral response related genes, but downregulation of inflammatory response related genes was observed (**Figures S5A and S5B**).

**Figure 5.**
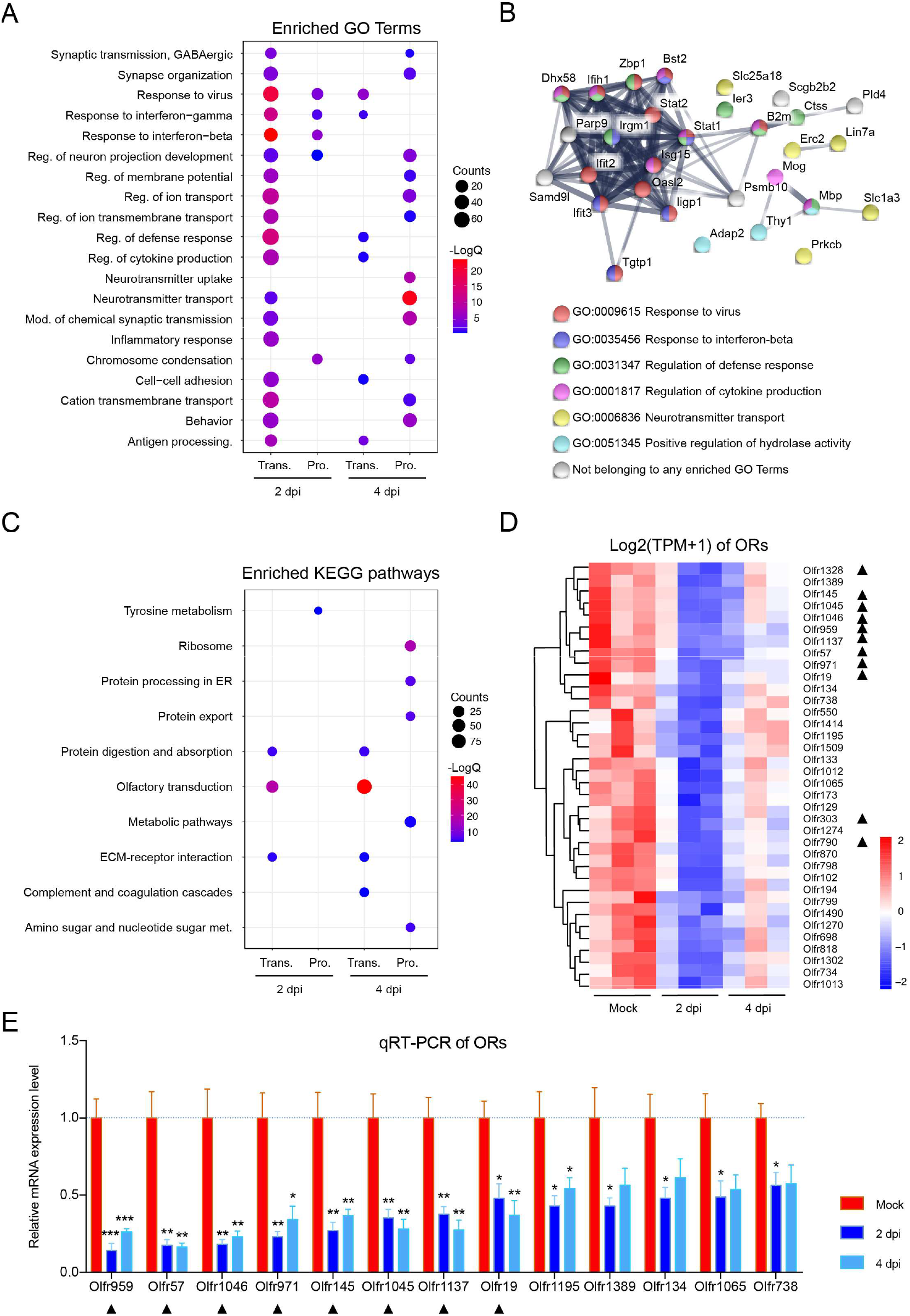
Host response to SARS-CoV-2 in OE at the mRNA and protein levels. (A) Dotplot visualization of enriched GO terms of up regulated genes/proteins at 2/4 dpi in OE. Gene enrichment analyses were performed using Metascape against the GO dataset for biological processes. “Reg.” for regulation, “mod.” for modulation, and “antigen processing.” for antigen processing and presentation of peptide antigen. (B) Interaction map of 30 proteins which consistently up regulated at both transcriptomic and proteomic levels along the course of SARS-CoV-2 infection in OE. Network nodes represent proteins, and their colors indicate different GO Terms they belonging to. Edges represent protein-protein associations, and their thickness indicates the strength of data support. (D) Dotplot visualization of enriched KEGG pathways of down regulated genes/proteins at 2/4 dpi in OE. Gene enrichment analyses were performed using String against the KEGG dataset. “Met.” for metabolism. The color of the dots represents the −LogQ value for each enriched KEGG pathways, and size represents the gene/protein counts enriched in each term. Heatmap indicating the expression patterns of 36 olfactory receptor genes which were significantly down regulated at 2 dpi. Colored bar represents Z-score of log_2_ transformed TPM+1. Total 11 of them also down regulated at 4 dpi were marked with black triangles. (E) RNA expression of 13 representative ORs by qRT-PCR. Columns with *, **, *** indicate ORs significantly down regulated at *p*<0.05, *p*<0.01 or *p*<0.001 relative to their Mock groups (One-way ANOVA followed by post hoc analysis with Turkey test, n = 3) respectively. Black triangles marked ORs were down regulated at both 2 and 4 dpi based on transcriptome data. See also Figure S3, S4 and S5. Table S1.

Of particular note, KEGG pathway enrichment of down regulated transcripts and proteins in OE showed that genes belonging to “olfactory transduction” were significantly enriched (**Figure 5C**). Among all 100 down regulated transcripts at 2 dpi, 36 were ORs (**Figures 5D and S3B**), while among 278 down regulated transcripts at 4 dpi, 97 were ORs (**Figures S3B and S4E**). Further RT-qPCR assay showed a dozen of OR genes were significantly down regulated in response to SARS-CoV-2 infection (**Figure 5E**), which may also attribute to the observed olfactory dysfunction.

## Discussion

In the present study, we used an established mouse model to demonstrate that SARS-CoV-2 infection can cause olfactory dysfunction and anosmia, and these experimental evidence support the hypothesis that SARS-CoV-2 infection as the cause of olfactory dysfunction and anosmia in COVID-19 patients (**Iravani et al., 2020**; **Moein et al., 2020**). The SARS-CoV-2 infected mice exhibited damaged OE, immune cell infiltration, down regulated OR expressions and impaired olfactory function, largely mimicking the olfactory abnormalities of COVID-19 patients. Robust viral replication and direct antiviral responses were detected in the OE of the infected mice, but not in OB and other parts of the brain, indicating that SARS-CoV-2 infection may be restricted in OM, instead of spreading to the CNS. A recent study also supports this point of view, for that SARS-CoV-2 protein can be detected in OE, but not in OB, in a hamster model (**Bryche et al., 2020**). One possible explanation for the absence of SARS-CoV-2 in CNS is the IFN-dependent antiviral mechanism, which is an effective barrier to limit the virus from invading into CNS (**Forrester et al., 2018**). In addition, the apoptosis of infected OSNs may contribute to the prevention of virus spreading into CNS after the rapid infection and destruction of OE (**Mori et al., 2002**).

Our results show that SARS-CoV-2 initially infects non-neuroepithelial cells, including sustentacular cells, Bowman’s gland cells and microvillar cells, which are involved in OSN support, host immune response, electrolyte balance maintenance, and mucus secretion (**Cooper et al., 2020**). Meanwhile, we observed various levels of damage in OE after SARS-CoV-2 infection, including cilia desquamating, loss of surface microvilli and substantial structural disorganization. In addition, our results showed a certain degree of cell apoptosis and inflammatory infiltration at both cell and molecular level following SARS-CoV-2 infection. All these data indicate that the damaged supporting non-neuroepithelial cells and inflammatory infiltration caused by SARS-CoV-2 infection contribute to the detrimental effects of the virus on olfactory function. Our results are supported by recent findings in mouse and human, showing that the non-neuroepithelial cells of OE express high levels of ACE2 and TMPRSS2 at both mRNA and protein levels (**Brann et al., 2020**; **Torabi et al., 2020**), (**Trotier et al., 2007**). Interestingly, SARS-CoV-2 positive signals were also observed in mOSNs and HBCs of infected animals, although we didn’t detect any hACE2 expression in these cells. The underlying mechanism remains elusive and a hACE2-independent spread of SARS-CoV-2 infection may be considered.

We observed many ORs were significantly down regulated at 2 and 4 dpi, suggesting the declined olfaction after SARS-CoV-2 infection. A recent study also showed that induction of anti-viral type I interferon signaling in the mouse OE was associated with diminished odor discrimination and decreased RNA levels of ORs (**Rodriguez et al., 2020**). These findings may support what we observed here that SARS-CoV-2 infection causes significant interferon response and dramatic OR decrease simultaneously in OE. We also observed three odorant-binding proteins (OBPs) significantly decreased at protein level with the infection of OE, which are compact globular water-soluble proteins with ligand-binding capabilities and thought to aid in capture and transport of odorants to the ORs (**Matarazzo et al., 2002**; **Pes and Pelosi, 1995**; **Sun et al., 2018**). Besides, although no virus infection was observed in OB, we detected some up/down regulated transcripts or proteins by transcriptomic and proteomic analyses. It worth noting that among all 4 proteins co-regulated at both transcriptomic and proteomic levels, Rtp1 (Receptor-transporting protein 1) was down regulated at both levels (Table S1). This protein specifically promotes functional cell surface expression of ORs (**Wu et al., 2012**), suggesting that the inhibition of Rtp1 in OB may lead to down regulation of ORs. Therefore, the damage of OE which is closely related to olfactory dysfunction are caused by SARS-CoV-2 infection of non-neuroepithelial cells and OSNs synergizes with the host antiviral immune responses.

According to our results, the olfactory dysfunction in SARS-CoV-2 infected animals is recoverable as almost all animals recovered to normal sense of smell at 4 dpi. Additionally, studies focusing on COVID-19 patients with anosmia has shown that most of them would recovery from loss of smell within a few weeks or less (**Hopkins et al., 2020**; **Yan et al., 2020**), indicating a potential mechanism of OE regeneration from injuries. The OE undergoes a lifelong regeneration and replacement depending on two populations of basal stem cells, HBCs and GBCs. HBCs are mitotically quiescent under the normal conditions and convert to be activated and differentiate into other kinds of cells once the damage of OE occurs (**Salazar et al., 2019**). Unlike HBCs, most of GBCs are mitotically activated and responsible for the regeneration of both neuronal and non-neuronal cells (**Gadye et al., 2017**; **Leung et al., 2007**; **Yu and Wu, 2017**). Indeed, we observed the regeneration of OE by the significant proliferation and morphological change of HBCs, accompanied by the differentiation of stem cells into iOSNs, sustentacular cells as well as microvillar cells. In this way, the structural basis and function of OE as well as the olfactory function can be restored to normal in SARS-CoV-2 infected animals. Furthermore, it was indicated that the damage and apoptosis of OSNs are closely involved in their regeneration (**Ishimura et al., 2008**), and the occurrence of inflammatory response also facilitates the stem cell differentiation and OE regeneration (**Chen et al., 2019**; **Lane et al., 2014**). At transcriptomic and proteomic level, we observed up regulated “regulation of neuron projection development” genes/proteins on 2 and 4 dpi, implying the progression of a neuron projection over time from its formation to the mature structure. Interestingly, although there were many significantly down regulated ORs on 4 dpi, the mRNA levels of many ORs rose back slightly compared with that of 2 dpi, indicating the OR expression tends to recover to normal.

In summary, our study established a mouse model of olfactory dysfunction induced by SARS-CoV-2. Considering the interspecies discrepancy of olfactory construction between rodent and human, e.g., the relative size of the OB to the brain, the proportion of the brain involved in olfaction and the expression of ORs (**Salazar et al., 2019**), further studies are recommended to reproduce the SARS-CoV-2 caused olfactory dysfunction in other animal models, especially the non-human primates. Also, to validate the targets and biological effects of SARS-CoV-2 infection in human specimens is still ponderable. The animal model of olfactory disorders is available to subsequently evaluate the antiviral drugs as well as vaccines for the inhibition of SARS-CoV-2 and the improvement of post viral olfactory disorders.

## Acknowledgments

We thank Drs. Changfa Fan, Jianfeng Liu, Bin Fu for critical reagents and helpful discussion. This work was supported by the National Key Research and Development Project of China (2016YFD0500304, 2020YFC0842200, and 2020YFA0707801). C.F.Q. was supported by the National Science Fund for Distinguished Young Scholar (No. 81925025), and the Innovative Research Group (No. 81621005) from the NSFC, and the Innovation Fund for Medical Sciences (No.2019RU040) from the Chinese Academy of Medical Sciences (CAMS). P.X. was supported by the CAMS Innovation Fund for Medical Sciences (No.2019RU006). J.Z. was supported by Youth Program of National Natural Science Foundation of China (No.82002148) from NSFC, and the China Postdoctoral Science Fund (No. 2020T130134ZX). R.T.L was supported by the China Postdoctoral Science Fund (No.2019M664012 and No. 2020T130135ZX).

## Author Contributions

C.F.Q., Q.Y., J.Z., and G.Y. conceived the project and designed the experiments. Q.Y., J.Z., G.Y. and Q.H. performed the majority of the experiments and analyzed the data; R.T.L., Y.Z., Q.C., R.R.Z., H.Q., Y.Q.D., X.F.L., S.J.W., J.H.S., H.M.Z., and T.Z. contributed specific experiments and data analysis. Y.Z. and P.X. contributed to proteomic analysis. C.F.Q., Q.Y., J.Z., G.Y., and R.T.L. wrote the manuscript with all the input from all authors. C.F.Q, X.Y., and P.X. supervised the study. All authors read and approved the contents of the manuscript.

## METHODS

### Cell and Virus

The Vero cells were maintained at 37℃ under 5% CO_2_ in Dulbecco’s modified Eagle essential medium (DMEM) supplemented with 10% heat-inactivated fetal bovine serum (FBS, Gibco), 10 mM HEPES and 1% penicillin/streptomycin. The SARS-CoV-2 strain BetaCoV/Beijing/IMEBJ05/2020 (Nos. GWHACBB01000000) was originally isolated from a COVID-19 patient. For virus propagation, Vero cells were incubated with SARS-CoV-2 and the culture supernatants were collected at 3 dpi. The stock of SARS-CoV-2 was serially diluted and titered on monolayers of Vero cells. Studies with infectious SARS-CoV-2 were conducted under biosafety level 3 (BSL3) facilities at the Beijing Institute of Microbiology and Epidemiology, AMMS.

### SARS-CoV-2 infection of hACE2 mice

The animal operation procedure was reviewed and approved by the Laboratory Animal Center, AMMS (approval number: IACUC-DWZX-2020-001). For intranasal infection, 5.4 × 10^5^ PFU of SARS-CoV-2 was instilled into the nasal cavity of 6-8 weeks old hACE2 mice anaesthetized with sodium pentobarbital at a dose of 50 mg/kg by intraperitoneal route. Mice were monitored daily and euthanized at 2 or 4 dpi to isolate tissues.

### RNA Extraction and real-time quantitative PCR

Quantification of SARS-CoV-2 RNA, hACE2 and OR mRNA transcript levels were performed by real-time quantitative PCR (RT-qPCR). Total RNAs were isolated using TRIzol reagent (Invitrogen, Carlsbad, CA, USA) according to the manufacturer’s instructions. SARS-CoV-2 RNA was measured with the primer-probe set: CoV-F3 (5’-TCCTGGTGATTCTTCTTCAGGT-3’), CoV-R3 (5’-TCTGAGAGAGGGTC AAGTGC-3’) and CoV-P3 (5’-AGCTGCAGCACCAGCTGTCCA-3’). The relative expression of hACE2 mRNA was measured with the primer set: ehACE2 F1 (5’-CGAAGCCGAAGACCTGTTCTA-3’) and ehACE2 R1 (5’-GGGCAAGTGTGG ACTGTTCC-3’). The expression of glyceraldehyde-3-phosphate dehydrogenase (GAPDH) served as the endogenous control, and the following primer set was used: 5′-CCAACCGCGAGAAGATGA-3′ and 5′-CCAGAGGCGTACAGGGATAG-3′. Amplification was performed using a One Step PrimeScript RT-PCR Kit (Takara Bio, Otsu, Japan), and the following real-time PCR conditions were applied: 42 ℃ for 5 min and 95 ℃ for 10 s followed by 40 cycles of 95℃ for 5 s and 60 ℃ for 20 s on an LightCycler^®^ 480 Instrument (Roche Diagnostics Ltd, Rotkreuz, Switzerland). The absolute quantification of SARS-CoV-2 RNA levels was performed by comparison to a standard curve and shown as SARS-CoV-2 RNA copies per mouse. The relative expression of hACE2 and OR mRNA levels was calculated according to the 2^−ΔΔCt^ method. Each sample was assayed with three repeats.

### The buried food pellet test (BFPT)

The standard BFPT was used to evaluate the olfactory function of SARS-CoV-2-infected mice and DMEM-treated mice as previously described (**Lehmkuhl et al., 2014**; **Yang and Crawley, 2009**). Mice were food-restricted to 0.2 g chow per day for 2 days before test and during the experimental period to ensure motivation. The food pellet was buried 1 centimeter below the surface of 3-centermeter-high bedding in a clear test cage (45 cm L× 24 cm W × 20 cm H). One mouse was placed in the center of the cage, and the latency for the mouse to uncover the pellet was recorded. The latency was defined as 300 seconds for the mouse which cannot find the pellet within 5 minutes.

### RNAscope in situ hybridization

RNAscope *in situ* hybridization (ISH) for SARS-CoV-2 RNA was performed with the RNAscope assay (Advanced Cell Diagnostics, Newark, CA, USA) according to the manufacturer’s instructions. Briefly, the tissues were isolated immediately after euthanasia and fixed in 4% paraformaldehyde (PFA) for 24 hours, and embedded in paraffin after being decalcified using the 10% EDTA solution, 4-μm-thick formalin-fixed paraffin-embedded (FFPE) slides were warmed at 60 ℃ for 1 h before they deparaffinized in xylene, rehydrated in a series of graded alcohols and pretreated with RNAscope target retrieval at 95 ℃. Slides were detected in situ using 2.5 HD Reagent Kit (BROWN) (Cat: 322310) and sense probe from the RNAscope ISH probe-V-nCoV2019-S (Cat: 848561) at 40 ℃ in HybEZ hybridization oven and then counterstained with hematoxylin.

### Multiplex immunofluorescent staining

The 4-μm-thick paraffin sections were deparaffinized in xylene and rehydrated in a series of graded alcohols. Antigen retrievals were performed in citrate buffer (pH=6) with a microwave (Sharp, R-331ZX) for 20 min at 95°C followed by a 20 min cool down at room temperature. Multiplex fluorescence labeling was performed using TSA-dendron-fluorophores (NEON 9-color All round Discovery Kit for FFPE, Histova Biotechnology, NEFP950). Briefly, endogenous peroxidase was quenched in 3% H_2_O_2_ for 20 min, followed by blocking reagent for 30 min at room temperature. Primary antibody was incubated for 2-4 h in a humidified chamber at 37°C, followed by detection using the HRP-conjugated secondary antibody and TSA-dendron-fluorophores. Afterwards, the primary and secondary antibodies were thoroughly eliminated by heating the slides in retrieval/elution buffer (Abcracker®, Histova Biotechnology, ABCFR5L) for 10 s at 95°C using microwave. In a serial fashion, each antigen was labeled by distinct fluorophores. Multiplex antibody panels applied in this study were: hACE2 (Abcam, ab108209, 1:200); tdTomato (Rockland, 600-401-379, 1:500); SARS-CoV-2 nucleocapsid protein (Sinobiological, 40143-R004, 1:1000); GAP43 (Abcam, ab75810, 1:1000); OMP (Abcam, ab183947, 1:1500); CK5 (Abcam, ab52635, 1:800); CK8 (Abcam, ab53280, 1:800); Sox9 (Abcam, ab185230, 1:500); Sox2 (CST, 23064, :400); CD73 (CST, 13160, 1:500); Furin (Abcam, ab108209, 1:400); Tmprss2 (Abcam, ab92323, 1:500); CD3 (CST, 78588, 1:300); CD8 (CST, 98941, 1:300); Cleaved caspase-3 (CST, 9664, 1:1000); CD103 (Abcam, ab224202, 1:300); Ly-6G (CST, 87048, 1:400); CD68 (CST, 97778, 1:300); and Granzyme B (Abcam, ab255598, 1:300). After all the antibodies were detected sequentially, the slices were imaged using the confocal laser scanning microscopy platform Zeiss LSM880.

### Histopathological analysis

The structural integrity of the mouse OE was analyzed using hematoxylin and eosin (H&E) staining according to standard procedures. Briefly, after being rehydrated in series of graded alcohols, the 4-μm-thick slides of mouse OE were stained with hematoxylin for 30 s and washed in water. Slides were then stained in eosin for 15 s and washed again in water.

### RNA library construction and sequencing

hACE2 transgenic mice before or after SARS-CoV-2 infection (2 or 4 dpi) as previously described were used for RNA-Seq. Total RNA from OE and OB were extracted using TRIzol (Invitrogen, Carlsbad, CA, USA) and DNase I (NEB, USA) treated, respectively. Sequencing libraries were generated using NEBNext® UltraTM RNA Library Prep Kit for Illumina® (<E7530L, NEB, USA) following the manufacturer’s recommendations and index codes were added to attribute sequences to each sample. The clustering of the index-coded samples was performed on a cBot cluster generation system using HiSeq PE Cluster Kit v4-cBot-HS (Illumina, San Diego, California, USA) according to the manufacturer’s instructions. After cluster generation, the libraries were sequenced on Illumina Novaseq6000 platform and 150 bp paired-end reads were generated. After sequencing, perl script was used to filter the original data (Raw Data) to clean reads by removing contaminated reads for adapters and low-quality reads. Clean reads were aligned to the mouse genome (Mus_musculus.GRCm38.99) using Hisat2 v2.1.0. The number of reads mapped to each gene in each sample was counted by HTSeq v0.6.0 and TPM (Transcripts Per Kilobase of exon model per Million mapped reads) was then calculated to estimate the expression level of genes in each sample.

### Large-scale proteome sample preparation and Tandem Mass Tags (TMT) labeling

The OE and OB tissues were disrupted by a Grinding Mill for six cycles of 5 s each with lysis buffer [9 M Urea, 10 mM Tris-HCl (pH 8.0), 30 mM NaCl, 10 mM iodoacetamide (IAA), 5 mM Na_4_P_2_O_7_, 100 mM Na_2_HPO_4_ (pH 8.0), 1 mM NaF, 1 mM Na_3_VO_4_, 1 mM sodium glycerophophate, 1% phosphatase inhibitor cocktail 2 (Sigma, St. Louis, USA), 1% phosphatase inhibitor cocktail 3 (Sigma, St. Louis, USA), and 1 tablet of EDTA-free protease inhibitor cocktail (Roche, Basel, Switzerland) for every 10 mL of lysis buffer] and 2 mm steel balls, respectively. The supernatants were obtained after centrifuging at 8,000 rpm for 10 min at 4°C. The protein lysates were inactivated at 56 ℃ for 30 min then stored at −80°C before further processing. Protein concentration was measured by a short Coomassie blue stained 10% SDS-PAGE as described (**Xu et al., 2009**). The same amount of protein (130 μg) from each sample was reduced with 5 mM of dithiothreitol (DTT), alkylated with 20 mM of IAA, precleaned with 10% SDS-PAGE (10%, 0.7 cm), and digested in-gel with a final concentration of 12.5 ng/μL for Ac-trypsin combined with endoproteinase lys-C provided by Enzyme & Spectrum (Beijing, China) with a ratio of 2:1 at 37°C for 12-14 h (**Zhao et al., 2016**; **Zhao et al., 2015**). The extracted peptides from OE and OB groups were labeled with TMT10 reagents according to the manufacturer’s instructions (Thermo Scientific, San Jose, CA, USA), respectively. Ten labeled channels were then quenched with 5% hydroxylamine and combined according to normalization value by the ratio checking. The mixed samples were vacuum dried.

### Peptide fractionation and LC-MS/MS analysis

The dried TMT labeled mixture were resuspended in 100 μL of buffer A [2% acetonitrile (ACN), pH10) and separated by a high pH reverse phase HPLC system (Rigol, L-3120, Beijing, China). The combined samples were injected into a Durashell C_18_ column (150 Å, 5 μm, 4.6 × 250 mm^2^) and eluted with a linear gradient in 60 min. Briefly, the solvent gradients of buffer B (2% dd H_2_O and 98% ACN) were as follows: 0% for 5 min, 0-3% for 3 min, 3-22% for 37 min, 22-32% for 10 min, 32-90% for 1 min, 90% for 2 min, and 100% for 2 min. The LC flow rate was set at 0.7 mL/min and monitored at 214 nm. The column oven was set at 45 °C. Total 60 fractions were collected and then combined into 15 fractions before vacuum drying according to the peak abundance. The combined samples were dissolved in loading buffer [1% ACN and 1% formic acid (FA)] and analyzed using an EASY-nLC 1200 ultra-performance liquid chromatography system (Thermo Fisher Scientific, San Jose, CA, USA) equipped with a self-packed capillary column (75 μm i.d. × 15 cm, 3 μm C_18_ reverse-phase fused-silica), with a 78 min nonlinear gradient at a flow rate of 600 nL/min. The gradient was comprised of an increase from 6% to 15% solvent B (0.1% FA in 80% ACN) for 15 min, 15% to 30% in 40 min, 30% to 40% in 15 min, 40% to 100% in 1 min, and finally holding at 100% for the last 7 min. The eluted peptides were analyzed on Orbitrap Fusion Lumos (Thermo Fisher Scientific, San Jose, CA, USA). MS_1_ data were collected in the Orbitrap using a 120 k resolution over an *m/z* range of 300-1500 setting the maximum injection time (MIT) to 50 ms. The automatic gain control (AGC) was set to 4×10^5^, determined charge states between 2 and 7 were subjected to fragmentation via higher energy collision-induced dissociation (HCD) with 37% collision energy and a 12 s dynamic exclusion window was used with isotopes excluded. For the MS/MS scans, the fractions were detected in the Orbitrap at a resolution of 50 k. For each scan, the isolation width was 1.6 *m/z*, the AGC was 5 × 10^4^, and the MIT was 86 ms.

### Database search

The raw files from OE and OB groups were searched with MaxQuant (v1.5.5.0) against the mouse reviewed proteome downloaded from UniProt containing 17,478 entries and a canonical SARS-CoV-2 proteome with 30 potentially viral proteins from the SARS-CoV-2 genome (NC_045512.2), and a common contaminant database (http://www.maxquant.org/contaminants.zip), respectively. Fully tryptic peptides with as many as 2 missed were allowed. TMT 10 plex (N-Term/K) and cysteine carbamidomethyl were set as fixed modification, whereas oxidation of methionine was set as variable modification. The tolerance of the precursor and fragment ions were set to 20 ppm.

### Bioinformatic analyses

DESeq2 v1.6.3 was used for differential gene expression analysis. Genes with padj≤0.05 and |Log_2_FC| > 1 were identified as differentially expressed genes (DEGs). The total proteome quantification datasets were median-normalized, and *pValue* was calculated by Perseus (1.6.6.0). Proteins ratios between control and infection ≥1.5-fold and *pValue* ≤ 0.05 were considered as regulated differentially expressed proteins (DEPs). DEGs and DEPs were used as query to search for enriched biological processes (Gene ontology BP) using Metascape (**Zhou et al., 2019**). KEGG pathway enrichment and protein interaction network were analyzed using STRING (**Szklarczyk et al., 2019**). Heatmaps of gene expression levels were constructed using pheatmap package in R (https://cran.rstudio.com/web/packages/pheatmap/index.html). Dot plots and volcano plots were constructed using ggplot2 (https://ggplot2.tidyverse.org/) package in R.

### Statistical analysis

Data were analyzed using GraphPad Prism 8 (GraphPad Software, San Diego, California, USA). The values shown in the graphs are presented as the mean ± standard deviation of at least three independent experiments. Statistical differences between groups were analyzed using two-tailed unpaired t-tests or a one-way ANOVA statistical test with Dunnett multiple comparisons tests; *p* < 0.05 was considered statistically significant.

