## Supplemental Information for "SARS-CoV-2 infection causes transient olfactory dysfunction in mice"

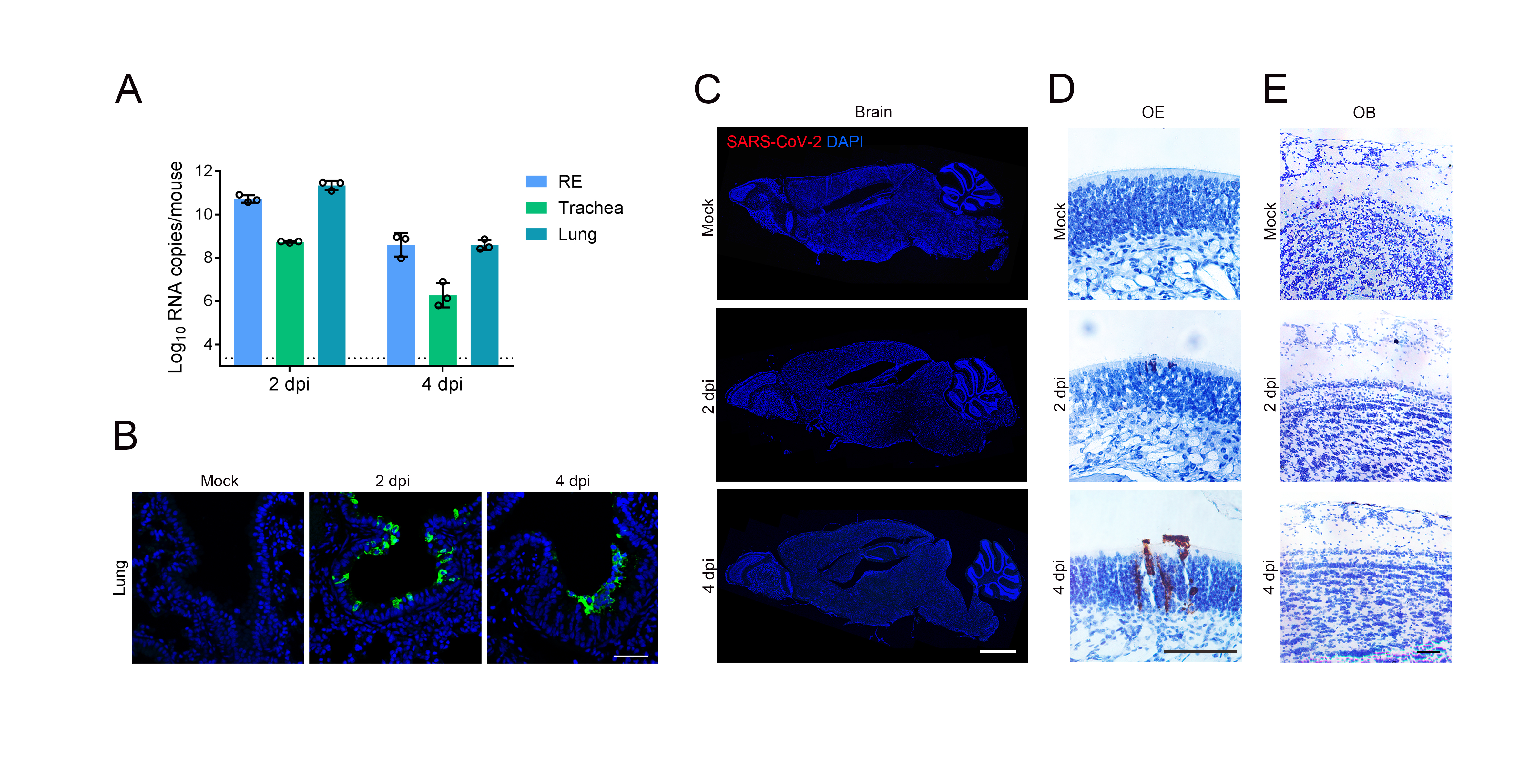

**Figure S1. SARS-CoV-2 infection in respiratory tract and olfactory system of hACE2 mice, related to Figure 1.**

(A) Viral RNA detection in nasal RE, trachea and lung of SARS-CoV-2 infected mice. Viral RNA copies were determined by real-time qPCR and shown as mean ± SD from three independent replicates.

(B) Representative immunostaining of lung tissue from SARS-CoV-2 infected or mock treated mice for SARS-CoV-2 N protein (green) and DAPI (blue). Scale bar, 50 μm.

(C) Representative immunostaining of brain tissues from SARS-CoV-2 infected and mock treated mice for SARS-CoV-2 N protein (green) and DAPI (blue). Scale bar, 2 mm.

(D and E) Representative RNAscope *in situ* hybridization (ISH) for SARS-CoV-2 RNA detection in OE (D) or OB (E) tissues from SARS-CoV-2 infected or mock treated mice. Scale bar, 100 μm.

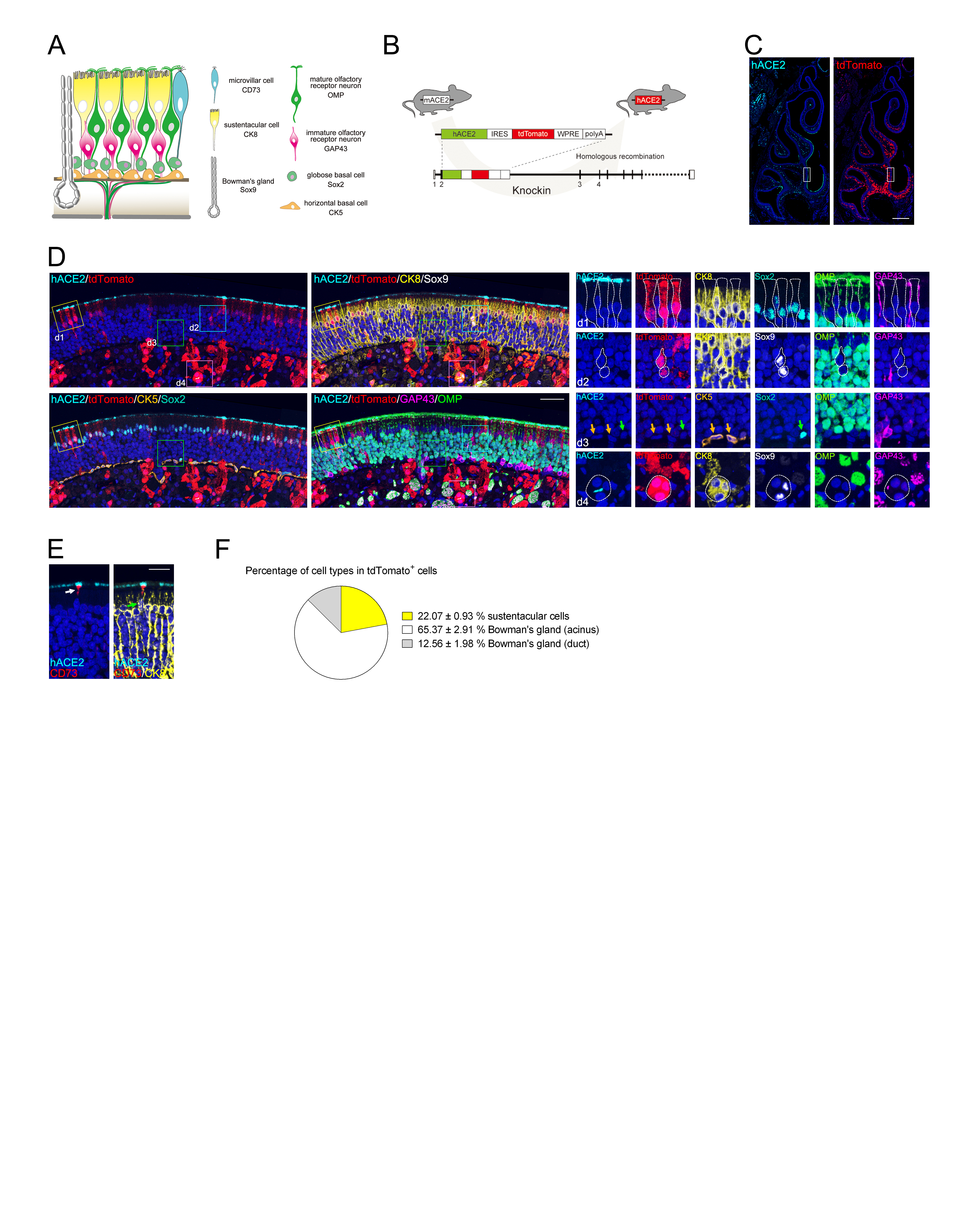

**Figure S2. hACE2 is mainly expressed by non-neuroepithelial cells in the OE of hACE2 mice****, related to Figure 2.**

(A) Schematic diagram shows the cell compartments of mouse OE and their representative markers.

(B) Schematic diagram shows the strategy of generating hACE2 mice by insertion of a hACE2-IRES-tdTomato cassette into the exon2 of endogenous mouse ACE2 gene.

(C) Representative immunofluorescent staining of hACE2 (cyan) and tdTomato (red) on coronal section of hACE2 mouse olfactory system. Scale bar, 500 μm.

(D) The framed area of (C) at larger magnifications with multiple cell markers including CK8 (yellow), Sox9 (white), CK5 (gold), Sox2 (emerald), GAP43 (magenta) and OMP (green). The framed areas in (D) are labeled as d1 (the surface region of OE), d2 (the duct of Bowman’s gland), d3 (the basal region of OE) and d4 (the acinus of Bowman’s gland), and are shown adjacently at larger magnifications with each marker separately displayed. The tdTomato+ cells were outlined by dashed lines. None of HBC or GBC expresses hACE2 or tdTomato (gold and green arrows, respectively). Scale bar, 50 μm.

(E) Representative multiplex immunofluorescent staining shows CD73/CK8-positive microvillar cells also express hACE2. Scale bar, 50 μm.

(F) Statistical analysis of the percentage of each cell compartment within the tdTomato-positive cells. Data were presented as mean ± SD (n = 3).

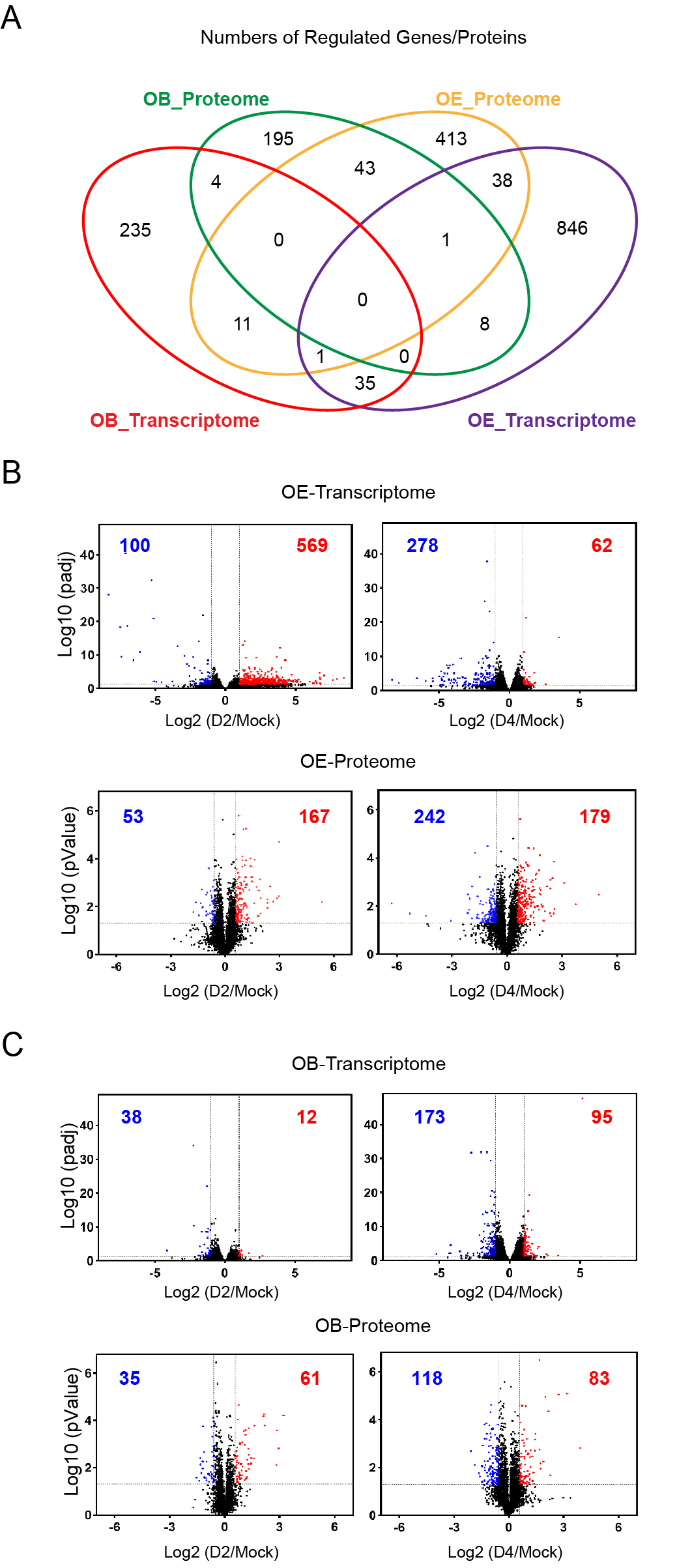

**Figure S3. The comparison of regulated genes or proteins corresponding to SARS-CoV-2 infection in OE and OB, related to Figure 5.**

(A) Comparison of regulated genes or proteins in SARS-CoV-2-infected OE and OB. The Venn diagram depicts genes or proteins shared and/or unique between each comparison.

(B) Volcano plots indicating differential regulated genes and proteins of OE along the course of SARS-CoV-2 infection.

(C) Volcano plots indicating differential regulated genes and proteins of OB along the course of SARS-CoV-2 infection. Up regulated genes (padj < 0.05) with a log_2_ (fold change) of more than 1 are indicated in red, down regulated genes (padj < 0.05) with a log_2_ (fold change) of less than -1 are indicated in blue. Up regulated proteins (*p-value* < 0.05) with a log_2_ (fold change) of more than 0.58 are indicated in red, down regulated proteins (*p-value* < 0.05) with a log_2_ (fold change) of less than -0.58 are indicated in blue.

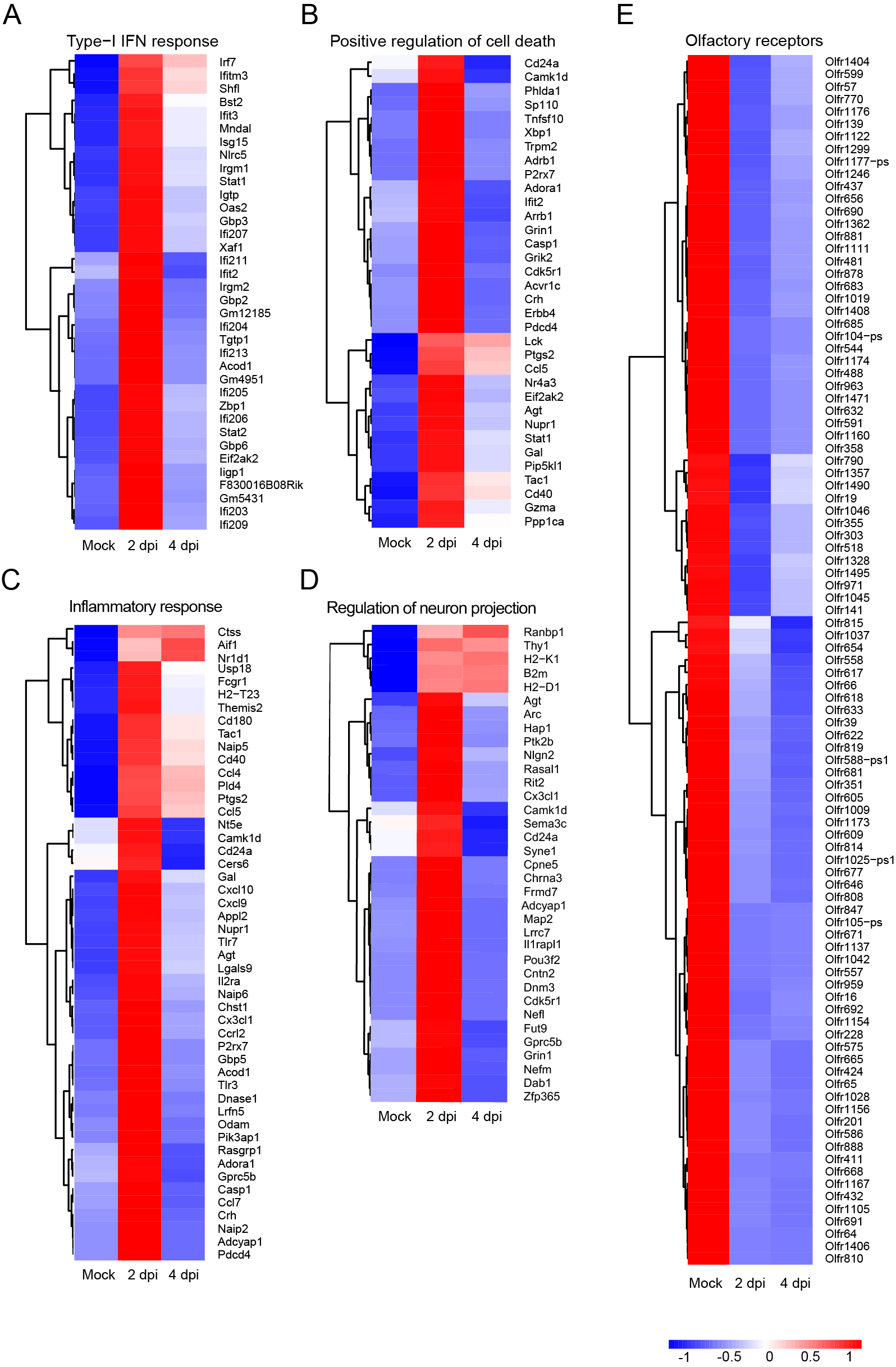

**Figure S4. Heatmaps depicting the expression levels of the regulated genes enriched in different GO annotating groups in OE, related to Figure 5.**

(A) Type I IFN response (GO:0035455 and GO:0035456).

(B) Positive regulation of cell death (GO:0010942).

(C) Inflammatory response (GO:0006954).

(D) Regulation of neuron differentiation (GO:0045664). The graphs depict the DEGs of infected compared with mock-treated OE at 2 and/or 4 dpi. Genes included have a | log_2_(fold change) | of more than 1 and a padj value of less than 0.05.

(E) Heatmap indicating the expression patterns of 97 olfactory receptor genes which were significantly down regulated at 4 dpi. Colored bar represents Z-score of log2 transformed TPM+1.

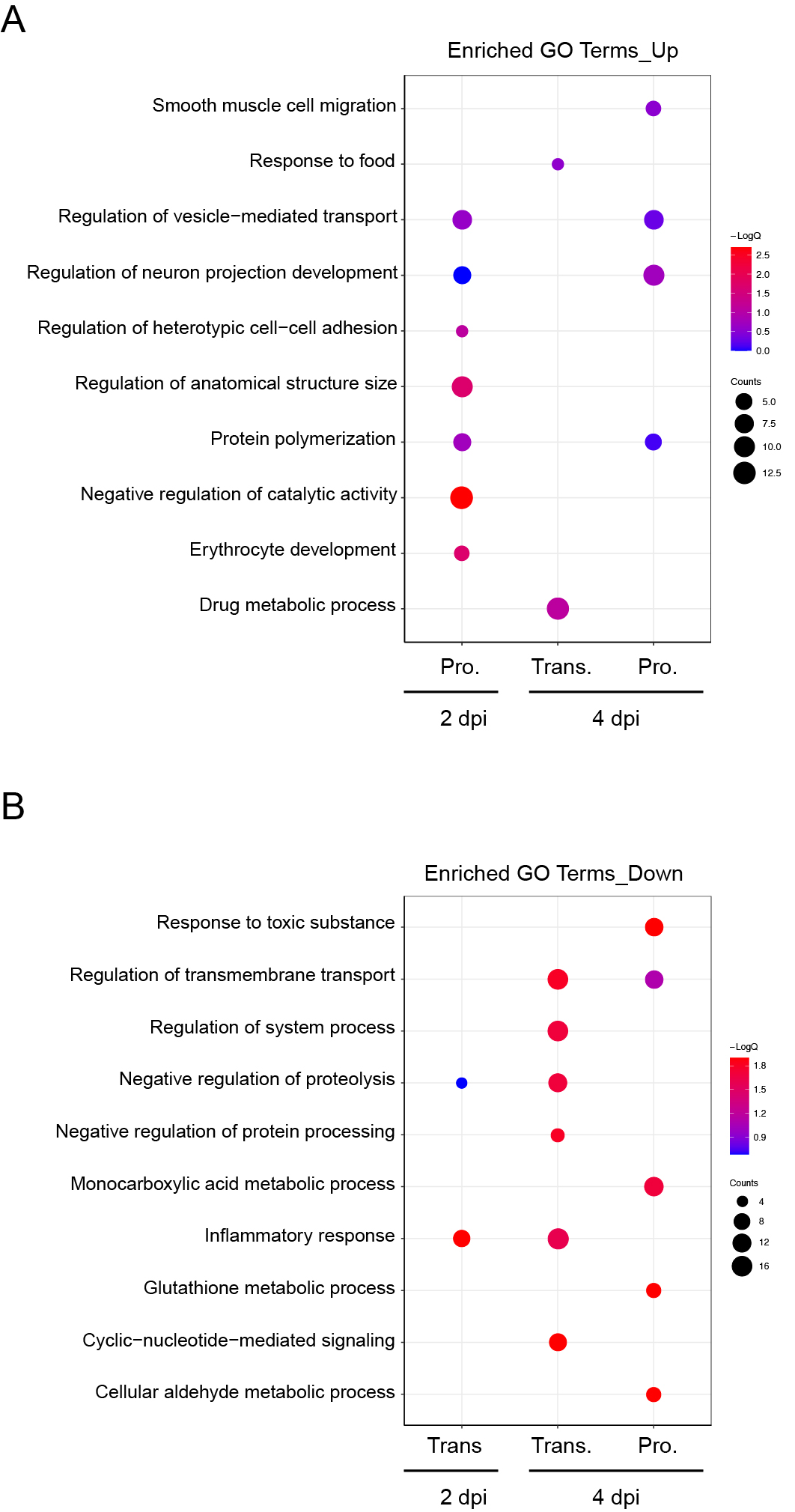

**Figure S5. Host response to SARS-CoV-2 in OB at mRNA and protein levels, related to Figure 5.**

(A) Dotplot visualization of enriched GO terms of up regulated genes/proteins on 2/4 dpi in OB. No GO terms were enriched using up regulated genes at 2 dpi.

(B) Dotplot visualization of enriched GO terms of down regulated genes/proteins at 2/4 dpi in OB. No GO terms were enriched using up regulated proteins at 2 dpi. Gene enrichment analyses were performed using Metascape against the GO dataset for biological processes.

**Table S1.** **Detailed information of 4 proteins co-regulated at both transcriptomic and proteomic level along the course of SARS-CoV-2 infection in OB, related to Figure 5.**

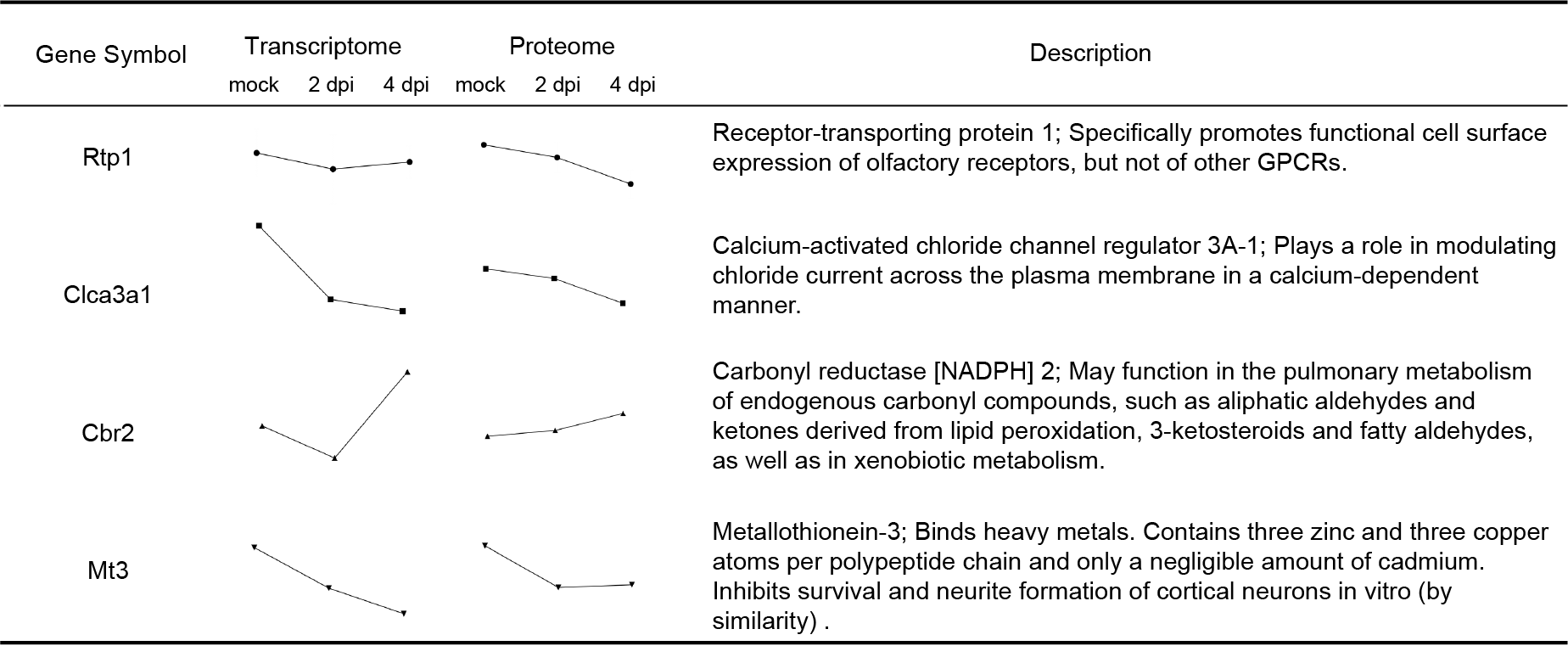
